# Macrophage Receptor with collagenous structure (MARCO) recognizes the cellular protein corona formed on environmental particles

**DOI:** 10.1101/2023.05.24.542216

**Authors:** Seishiro Hirano, Sanae Kanno

## Abstract

Several difficulties hamper the toxicological evaluation of particulate substances *in vitro*. One of the most serious difficulties is that most particles are insoluble in water, and easily agglomerate during the assay. Albumin and some detergents have been used to disperse the particles in the culture medium. However, effects of these additives on the cellular uptake of particles and the following biological consequences are unknown. The first event when phagocytic cells encounter particulate substances is an association of the plasma membrane with the particle surface. Macrophage receptor with collagenous structure (MARCO) has been identified as a receptor for environmental particles such as titanium dioxide (TiO_2_) and carbonaceous particles. However, it remained to be revealed how MARCO recognizes the unopsonized particles. While investigating the particle surface-cell membrane interaction using GFP-tagged MARCO-expressing cells, we found that cytosolic proteins were released from the cells and rapidly adsorbed on the surface of TiO_2_ particles. The GFP-tagged MARCO-overexpressing cells were further engineered to express RFP-tagged LC3 (cytosolic protein) to visualize adsorption of proteins on the surface of titanium dioxide (TiO_2_) particles. We found that cytosolic proteins including LC3 were released in the culture medium from intact cells and adsorbed on the particle surface quickly in the cell culture system. These results indicate that MARCO probably recognizes adsorbed intracellular proteins rather than the uncoated inorganic surface of environmental particles.

## Introduction

Macrophage receptor with a collagenous structure (MARCO) was first cloned and characterized as a receptor that binds to bacteria and acetylated-LDL in murine spleen and lymph nodes (Elomaa et al. 1995). MARCO (SR-A1/SR-A6) is inducible by LPS and plays an important role in the host antibacterial defense (van der Laan et al. 1999). Since MARCO was found to be a receptor of phagocytotic cells for unopsonized particulate substances (Kobzik 1995; Palecanda et al. 1999), a variety of substances such as carboxylate-modified polystyrene (Kanno et al. 2007), latex beads (Arredouani et al. 2005), titanium dioxide (TiO_2_) (Arredouani et al. 2005), and carbon nanotubes (Hirano et al. 2010; Hirano et al. 2008) have been reported to be recognized by MARCO. It should be noted that MARCO also mediates the cellular internalization of exosomes (Kanno et al. 2020), and phagocytosis of tumor cells by macrophages proceeds via a MARCO-integrin β_5_ interaction (Xing et al. 2021). MARCO is thought to be involved in the accumulation of PIP2 in the plasma membrane at the site of physical contact which is followed by recruitment of moesin, a cytoskeletal cross-linker protein (Mu et al. 2018). However, it is still uncertain what molecules or structures MARCO recognizes preferentially as ligands.

Binding of FcγR to IgG and that of complement receptor 3 (CR3, α_M_β_2_ integrin) to C3bi is the first step of opsonin-mediated phagocytosis of microorganisms or particles (Caron and Hall 1998; Jaumouille et al. 2019; Lam et al. 2009). In contrast, identification of ligands on environmental particles for MARCO-mediated phagocytosis is missing. Even though MARCO appears to recognize unopsonized environmental particles, the surface of the particles may not be “bare” and should be modified by substances in the culture medium and biological fluid *in cellulo* assay. Bovine serum albumin (BSA) is a major component of the protein corona formed on the nanoporous poly(methacrylic acid) polymer particles following exposure to FBS-containing media (Yan et al. 2013. SR-A, expressed on PMA-stimulated macrophage-like THP-1, recognized the conformationally changed BSA on the particles (Yan et al. 2013). In contrast to the stimulated THP-1 cells, the cell-to-bare particle association was decreased by BSA in unstimulated monocyte-like THP-1 (Lunov et al. 2011a). It is interesting to note that undifferentiated THP-1 cells internalize the particles through endocytosis, while PMA-differentiated THP-1 cells internalize the particles via macropinocytosis (Lunov et al. 2011a).

The protein corona formation changes the toxicity of environmental particles. The presence of a hard protein corona, tightly bound proteins on the surface, promoted cellular uptake of nano-silica particles in both A549 and RAW264.7 cells. However, the cytotoxicity of the particles was reduced by the corona, suggesting a possibility that the silica surface interacts with the cell membrane and perturbs membrane integrity (Leibe et al. 2019). Another report indicated that adherence of silica nanoparticles to A549 lung cells in serum-free culture medium was higher than that in the complete medium. In this study the cellular content of ATP was decreased by silica nanoparticles dose-dependently only when the cells were cultured in the absence of serum (Lesniak et al. 2012). It has also reported that graphene oxide-FBS protein binding reached an equilibrium within 30 min and the cytotoxicity of graphene oxide was reduced when the particles were coated with FBS proteins (Hu et al. 2011). Binding of MARCO to 100 nm silica particles is decreased by human serum in a concentration-dependent manner and further reduced by BSA, but not by well-known MARCO ligands such as LPS and acetylated LDL (Lara et al. 2018). These observations suggest that corona proteins which MARCO preferentially recognizes are different from albumin.

In addition to BSA in the culture medium, secretory proteins released from the cells are adsorbed on the particle surface. Bare silica nanoparticles adsorbed cytosolic, cytoskeleton, and cell membrane-associated proteins released in the serum-free culture medium in 1 h (Lesniak et al. 2012). ZnO nanowires adsorbed proteins released from human peripheral blood-derived macrophages grown in serum-free medium (Muller et al. 2010). Thus, it is important to investigate the protein corona formed by intracellular components in the cell culture system. In the present report, we report that cytosolic proteins are released rapidly from living cells and MARCO probably recognizes particulate substances via intracellular proteins adsorbed on the particle surface.

## Materials and Methods

### Chemicals and Particles

Wheat germ agglutinin (WGA)-AlexaFluor594 and pHrodo red dextran for endocytosis (10,000 MW, P35368) were obtained from Molecular Probe/ThermoFisher (Eugene, OR). Glutaraldehyde-stabilized sheep red blood cells (SRBC, InterCell Tech., Jupiter, FL) was used at a final concentration of 2.5 μg/mL. Dynasore (an inhibitor of macropinocytosis), 5-(N-ethyl-N-isopropyl) amiloride (EIPA, an inhibitor of endocytosis), and spherical TiO_2_ particles (s-TiO_2_, <100 nm, #637262) were obtained from Sigma-Aldrich (St. Louis, MO). The hydrodynamic characteristics of s-TiO_2_ particles were described elsewhere (Hirano and Kanno 2020). A sample of fibrous TiO_2_ (f-TiO_2_) particles was a gift from Japan Fibrous Material Research Association (TO1, JFMRA, Tokyo, Japan). The nominal average length and diameter of f-TiO_2_ particles are 2.1 and 0.14 μm, respectively. The TiO_2_ particles were heat-treated (250 °C, 2 h) in an electric furnace before use to remove any potentially adsorbed organic substances including endotoxin.

### Cells

Preparation methods of CHO-K1 and HEK293 cells stably expressing GFP-MARCO were described elsewhere (Hirano and Kanno 2015). These cells were designated as GFP-MARCO-CHO and GFP-MARCO-HEK, respectively. CHO-K1 cells stably expressing antioxidant responsive element (ARE) were used as negative control for GFP-MARCO-CHO, and was designated as ARE-CHO (Hirano et al. 2013). GFP-MARCO-CHO and GFP-MARCO-HEK cells were further transduced with RFP-LC3 plasmid which was a gift from Tamotsu Yoshimori (Addgene, plasmid #21075) (Kimura et al. 2007). The stable transfectants were selected using G418, and the cells expressing RFP-LC3 in GFP-MARCO-CHO and GFP-MARCO-HEK cells were designated as gMARCO-rLC3-CHO and gMARCO-rLC3-HEK, respectively. CHO- and HEK-derived cells were sub-cultured in F12 and DMEM media, respectively, containing 100 units/mL penicillin, 100 μg/mL streptomycin, and 10% heat-inactivated fetal bovine serum (FBS).

### Microscopic observation

The live gMARCO-rLC3-CHO and gMARCO-rLC3-HEK cells were observed by laser confocal microscopy (TCS-SP5, Leica Microsystems, Solms, Germany) or fluorescence microscopy (Eclipse TS100, Nikon, Tokyo, Japan). TiO_2_ particles treated with conditioned medium were also observed by laser confocal microscopy.

### Preparation of culture medium condensates and cell lysates

Pre-cultured cells in FBS-containing complete medium were rinsed once and further cultured in Opti-MEM I (Gibco/ThermoFisher, Grand Island, NY) to analyze proteins released in the culture medium. The culture medium was first centrifuged at 120 × *g* for 5 min at room temperature to remove cell debris and the supernatant was concentrated by 50 folds before SDS-PAGE electrophoresis. The concentration was performed by ultrafiltration (4 °C, 3200 × *g*, 100-120 min) using an Amicon Ultra (MW 3,000, Millipore, Cork, Ireland). The medium change to FBS-free Opti-MEM I was required to concentrate the culture medium by ultrafiltration because the viscosity of FBS hampered the ultrafiltration.

Cell monolayers were rinsed twice with PBS and lysed on ice for 10 min with cold RIPA lysis buffer (1× TBS, 1% Nonidet P-40, 0.5% sodium deoxycholate, and 0.1% SDS) containing protease inhibitors (Santa Cruz, Dallas, TX) and a phosphatase inhibitor cocktail (Pierce/ThermoFisher, Rockford. IL). Each lysate was transferred to a microcentrifuge tube and centrifuged at 9000 × *g* for 5 min at 4 °C.

### Immunoblot analysis

Aliquots of the concentrated culture medium and the supernatant of cell lysates were mixed with LDS sample buffer (1 × TBS, 10% glycerol, 0.015% EDTA, 50 mM DTT, and 2% LDS) and heated at 95 °C for 5 min. Proteins were separated on a 4-12% SDS-PAGE gel and then electroblotted onto a PVDF membrane. The membrane was blocked with PVDF blocking reagent (TOYOBO, Osaka, Japan) and probed with primary antibodies (1:500 dilution, 1 h) followed by HRP-tagged secondary antibodies (1:2000 dilution, 45 min). The antibodies were dissolved in Can-Get-Signal^®^ solution (TOYOBO). The membrane was soaked in ECL (Prime, GE Healthcare, Buckinghamshire, UK) and the chemiluminescence was detected by a CCD camera (Lumino Imaging Analyzer, FAS-1100, TOYOBO). The following antibodies were used for western blotting: anti-LC3 (8E10, MBL, Tokyo, Japan), anti-RFP cocktail (1G9-3G5, MBL), anti-BSA (1C11, Abcam, Cambridge, UK), anti-ERK (137F5, Cell Signaling, Danvers, MA), HRP-tagged anti-GAPDH (3H12, MBL), HRP-tagged tubulin (PM054, MBL), HRP-tagged goat anti-mouse IgG (sc-2354, Santa Cruz), and HRP-tagged goat anti-rabbit (Cell Signaling) antibodies.

### Statistical analyses

Data are presented as means ± SD or SEM. Student’s *t*-test was applied to compare the mean values between two groups. Probability values less than 0.05 were accepted as indicative of statistical significance.

## Results

### Plasma membrane activity of GFP-MARCO-CHO cells

First, we examined whether MARCO was involved in the association of cells with biological particles as well as environmental particles. Fig. 1 shows that the number of SRBC associated with cell bodies was significantly increased in GFP-MARCO-CHO than ARE-CHO cells. Some RBCs were tethered by the dendritic structures which are unique to MARCO-expressing cells (Kimura et al. 2007). These results show that MARCO recognized biomolecules and increased the adherence of cells to RBCs. The plasma membrane of CHO cells was completely internalized into endosomes/lysosomes in 30 min as WGA-bound plasma membranes (0 min) disappeared from the cell surface and co-localized with GFP-MARCO in peri-nuclear granules (Fig. 2). These results indicate that the plasma membrane receptor MARCO was continuously internalized and processed in lysosomes. The lysosomal accumulation of GFP-MARCO was confirmed by treating the cells with chloroquine, an alkalinization agent for lysosomes, on the basis that GFP fluorescence is partially quenched at lower pH (Kneen et al. 1998). As we expected, the green fluorescence was intensified by chloroquine in GFP-MARCO-CHO cells (Supplementary Fig. 1). Reciprocally, the red fluorescence was quenched by chloroquine since pHrodo red dextran elicits bright red fluorescent signal only in the pH range of 5 – 10.

**Fig. 1.**
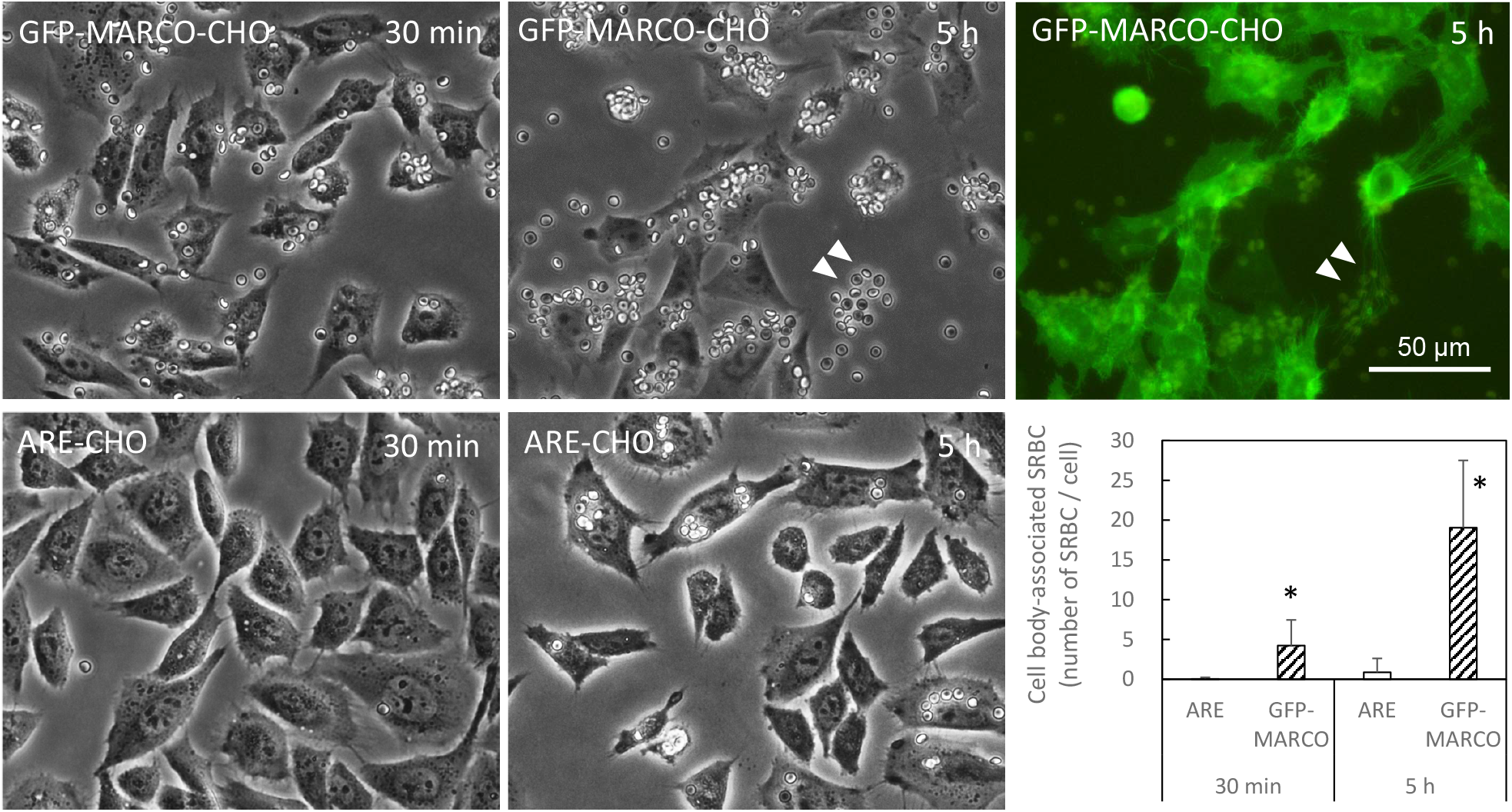
Cellular association of sheep red blood cells (SRBC) via MARCO receptors. GFP-MARCO-CHO or ARE-CHO (irrelevant vector-transfected) cells were incubated with glutaraldehyde-fixed SRBC at a concentration of 2.5 μg/mL for 30 min and 5 h in F12 complete medium containing 10% FBS. The cells were washed with HBSS 3 times. Bright filed and green fluorescence images of the live cells were captured by inverted fluorescence microscopy. Note that SRBC were also tethered by the dendritic structures formed on the bottom of the culture dish by GFP-MARCO-CHO cells (arrowheads). The number of SRBC associated with the cell body of GFP-MARCO-CHO cells was higher than that of ARE-CHO cells. Data are presented as mean ± SD. More than 100 cells were examined. See also the raw data file of Supplementary Table 1. *, Significantly different from the corresponding ARE-CHO cells.

**Fig. 2.**
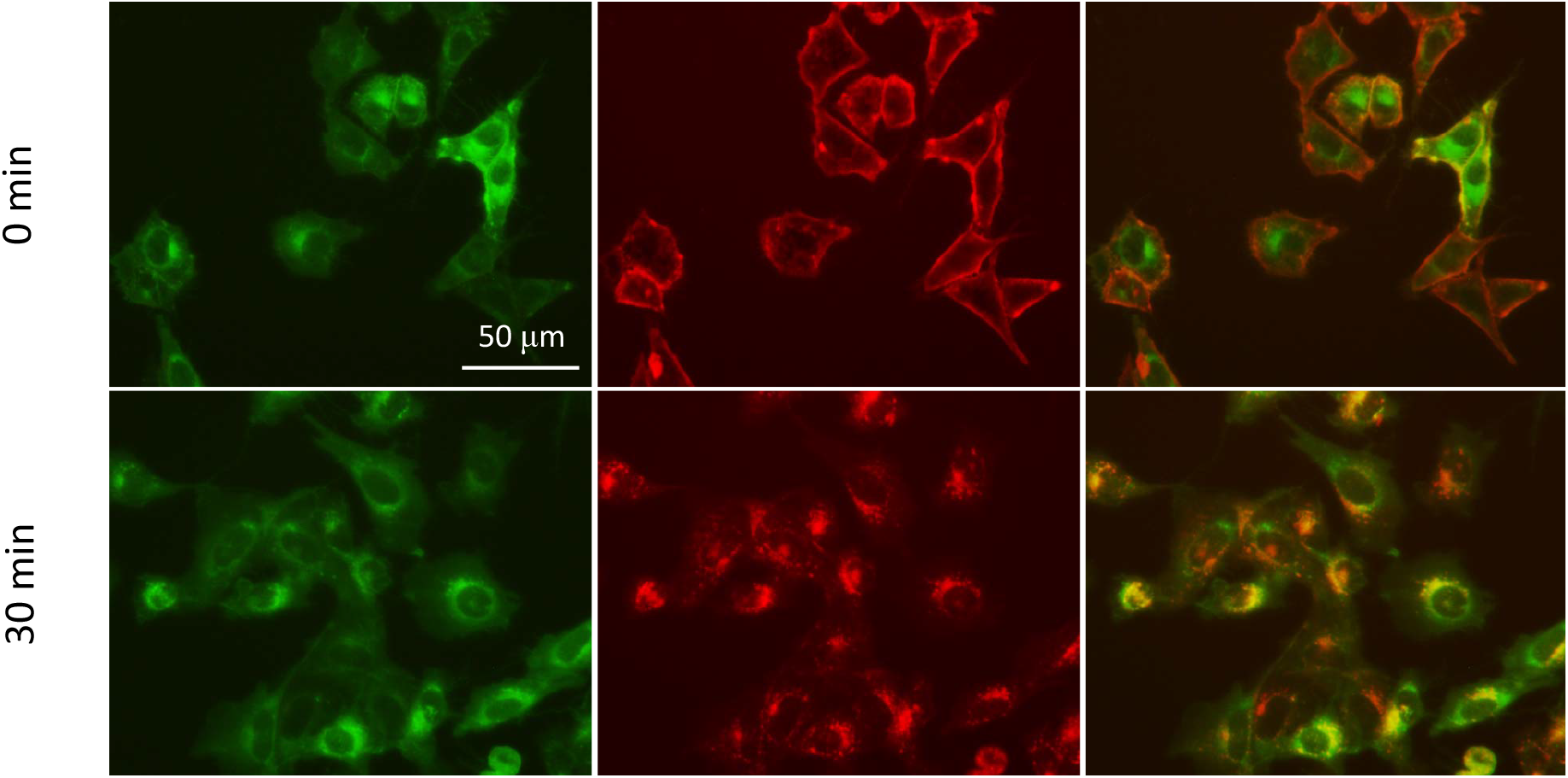
Internalization of plasma membrane in GFP-MARCO-CHO cells. The cells were labelled with 5 μg/mL wheat germ agglutinin (WGA)-Alexa Fluor® 594 Conjugate at 4ºC for 10 min. The cells were washed with cold HBSS three times and incubated in culture medium at 37ºC for 30 min. The green and red fluorescence images of the live cells were captured by inverted fluorescence microscopy. Note that WGA on the plasma membrane was completely internalized in 30 min and co-localized with GFP-MARCO at peri-nuclear regions.

### Release of intracellular proteins from gMARCO-rLC3-CHO cells

We used gMARCO-rLC3-CHO cells to differentiate between plasma membrane-bound (MARCO) and cytosolic proteins (LC3). Surprisingly, s-TiO_2_ and f-TiO_2_ particles were coated with RFP-LC3 before association with gMARCO-rLC3-CHO cells (Fig. 3A). During the particle internalization, RFP-labelled TiO_2_ appeared to be further coated with GFP-MARCO (arrowhead). Next, f-TiO_2_ particles were incubated with the supernatant of F12 conditioned medium containing 10% FBS to determine whether RFP-LC3 released into the culture medium was adsorbed on the particles. The confocal microscopic study showed that f-TiO_2_ particles were indeed coated with RFP-LC3 and GFP-MARCO small dots (0.5 – 2 μm) were observed sparsely on the fibers (Fig. 3B). These results indicated that cytosolic proteins were released from cultured cells and the proteins were adsorbed on the particle surface before being taken up by the cells. Then, f-TiO_2_ particles were incubated in fresh or conditioned 10% FBS-containing complete medium *in vitro*, and adsorbed proteins were analyzed by western blotting. BSA was detected when the particles were incubated in both fresh and conditioned media. RFP-LC3 was found to be adsorbed on the particle surface when the particles were incubated in the conditioned medium (Supplementary Fig. 2).

**Fig. 3.**
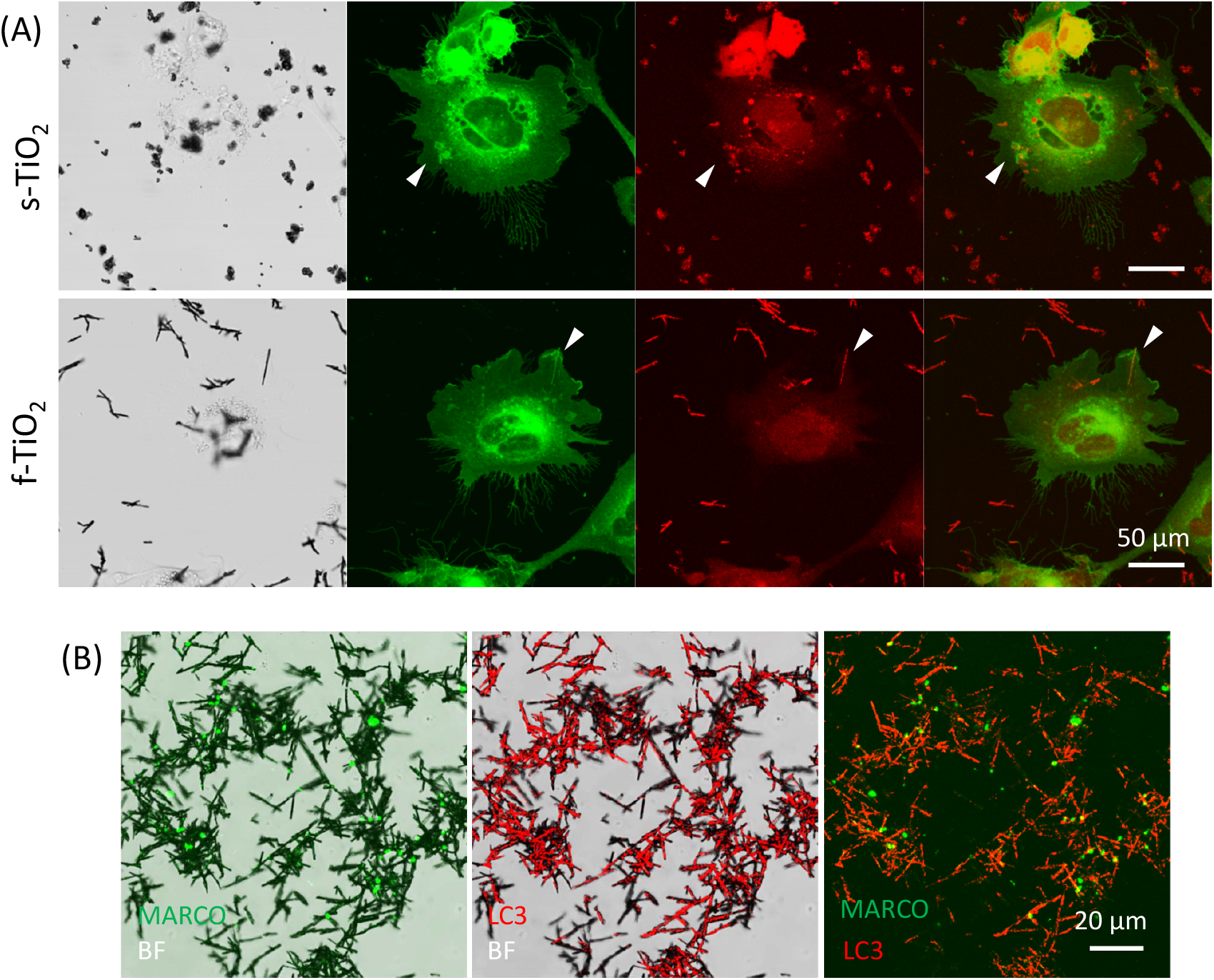
Cellular association of spherical (s-) and fibrous TiO_2_ (f-TiO_2_) particles with gMARCO-rLC3-CHO cells (A), and adsorption of RFP-LC3 on f-TiO_2_ in the cell-free conditioned medium (B). (A) The cells were cultured in 10% FBS containing complete F12 culture medium for 24 h, and then exposed to s-TiO_2_ or f-TiO_2_ at a concentration of 10 μg/mL for 1 h. The bright field and fluorescence images were captured by confocal microscopy. The arrowhead indicates internalized particles. (B) The conditioned medium was collected from sub-cultured cells in 10% FBS containing complete F12 culture medium. The sample was centrifuged at 150 × g for 5 min and the supernatant was dosed with f-TiO_2_ at a final concentration of 10 μg/mL and incubated at room temperature for 1 h with intermittent mixing. The f-TiO_2_ suspension was centrifuged at 9000 × g for 5 min. After removal of supernatant, f-TiO_2_ particles were re-suspended in PBS by pipetting, and a drop of the suspension was placed on a glass slide. The bright field and fluorescence images were captured by confocal microscopy.

Sub-confluent gMARCO-rLC3-CHO cells were cultured for up to 16 h in FBS-free Opti-MEM I culture medium, and cellular proteins released into the culture medium were detected by western blotting. It was found that 2 h of culture is enough to detect cytosolic proteins such as RFP-LC3 and GAPDH (Fig. 4A). Surprisingly, even though the culture time was reduced to 15 – 90 min, these cytosolic proteins were still detectable (Fig. 4B). ERK kinase, another cytosolic protein, was released into the culture medium in an analogous way, indicating that the cytosolic proteins were not released selectively, if any. The rapid release of the cytosolic proteins was temperature-dependent because these proteins were barely detected when the cells were incubated at 4 °C (Fig. 5). These results indicated that the cytosolic proteins were released from cultured cells during the normal cellular activity. Live gMARCO-rLC3-CHO cell images were captured by confocal microscopy with the time-lapse mode to find a possibility that the cytoplasmic components were leaked from live cells. We found the lamellipodia-like dendritic structures were extending and swelling at nods (Supplementary movie 1), suggesting that excess stress was imposed on the plasma membrane around the dendritic structure which may have rendered the membrane leaky for cytosolic proteins. However, it is not clear whether the stress was large enough for the leakiness of the plasma membrane.

**Fig. 4.**
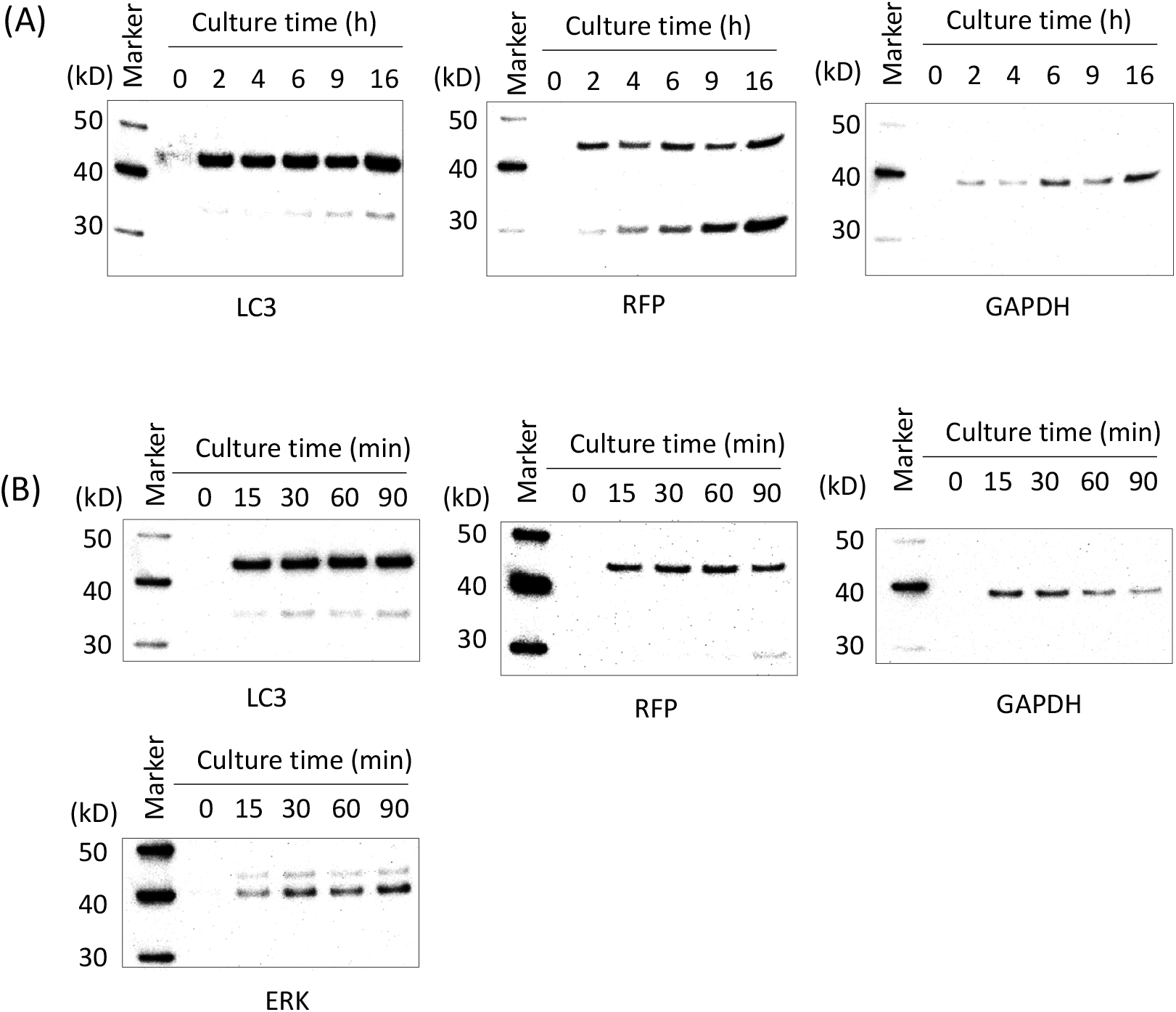
Release of cellular proteins from gMARCO-rLC3-CHO cells into culture medium. (A) gMARCO-rLC3-CHO cells were first grown in the complete F12 culture medium containing 10% FBS. The cells were rinsed with warmed FBS-free Opti-MEM I medium and further cultured in fresh Opti-MEM I medium for 2, 4, 6, 9, and 16 h. The culture supernatant was concentrated by 50 folds by ultrafiltration using an Amicon Ultra-4 centrifugal membrane (MW 3,000) for western blot analyses using anti-LC3, anti-RFP, and anti-GAPDH antibodies. (B) The culture medium was collected after 15, 30, 60, and 90 min of culture and concentrated by 50 folds. LC3, RFP, GAPDH, and ERK in the concentrated medium were detected by western blotting.

**Fig. 5.**
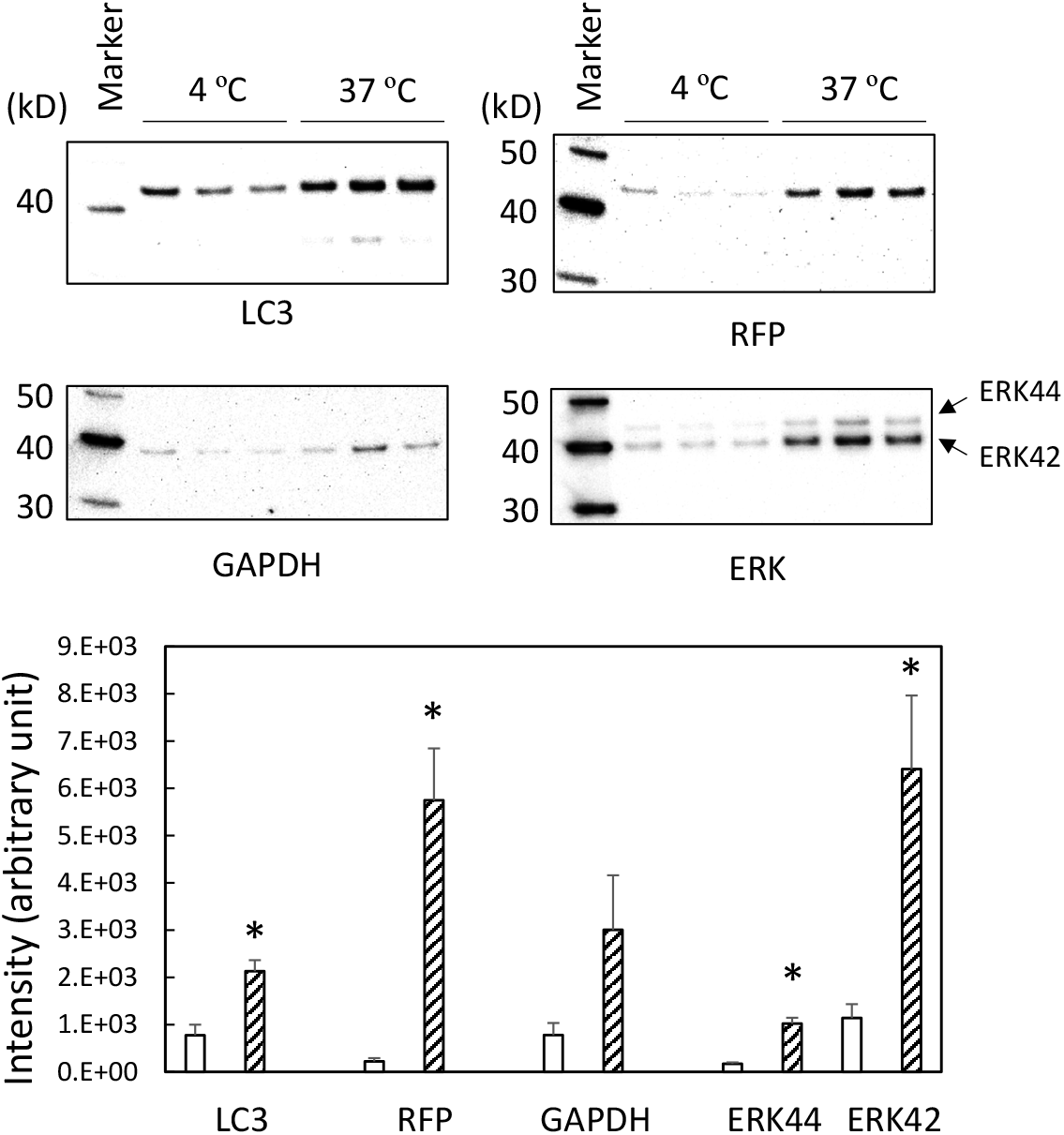
Temperature-dependent release of cellular proteins from gMARCO-rLC3-CHO cells into the culture medium. The cells were cultured in cold (4 ºC) HBSS or warmed (37 ºC) Opti-MEM I medium for 30 min in triplicate wells. Cytosolic proteins such as LC3, RFP, GAPDH, and ERK were detected by western blotting. See also the legend to Fig. 4. The abundance of proteins in the samples were estimated by densitometric analyses of the protein bands. Data are presented as mean ± SEM (N = 3). Open column, 4 ºC; Hatched column, 37 ºC. *, A significant difference was observed between 4 ºC and 37 ºC (p <0.05).

### Release of intracellular proteins in gMARCO-rLC3-HEK cells

The rapid release of cytosolic proteins was further confirmed using gMARCO-rLC3-HEK, HEK293 cells stably expressing both GFP-MARCO and RFP-LC3. The excess RFP-LC3 looked agglomerated in the cytoplasm of gMARCO-rLC3-HEK cells. gMARCO-rLC3-HEK cells formed faint dendritic structures on the glass bottom although they were much thinner than those of gMARCO-rLC3 CHO cells (Fig. 6A). The cytosolic proteins were released into the culture medium from gMARCO-rLC3-HEK cells in a time-dependent manner (Fig. 6B). The release of proteins in gMARCO-rLC3-HEK cells, however, appeared to be slower than that in gMARCO-rLC3-CHO cells (Figs. 4 and 6B).

**Fig. 6.**
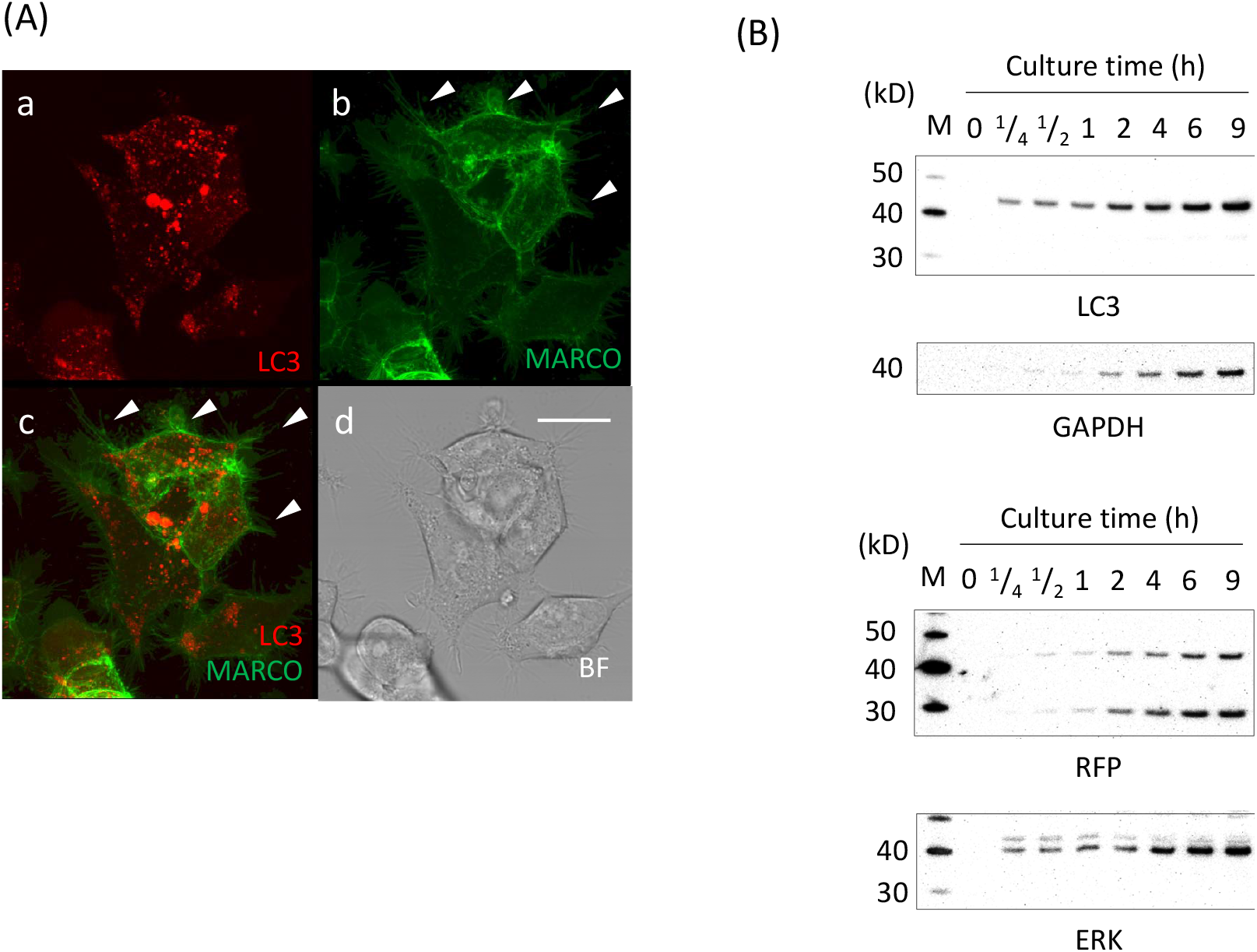
Confocal microscopic observation of gMARCO-rLC3-HEK cells, and release of cellular proteins into culture medium from gMARCO-rLC3-HEK cells. (A) The cells were cultured in complete DMEM culture medium containing 10% FBS. The arrowhead indicates the faint dendritic structures. (B) The cell monolayer was rinsed with warmed FBS-free Opti-MEM I medium and further cultured in fresh Opti-MEM I medium for up to 9 h. The cell-free conditioned medium was concentrated by 50 folds by ultrafiltration using an Amicon Ultra (MW 3,000) centrifugal filter for western blot analyses using anti-LC3, anti-RFP, anti-GAPDH, and anti-ERK antibodies.

### Effects of macropinocytosis and endocytosis inhibitors on the release of cytosolic proteins

MARCO-rLC3-CHO and gMARCO-rLC3-HEK cells were treated with EIPA and dynasore and the release of RFP-LC3 were measured. The release of RFP-LC3 was not affected by these inhibitors (Fig. 7), suggesting that inhibition of neither endocytosis nor macropinocytosis changed the release of cytosolic proteins.

**Fig. 7.**
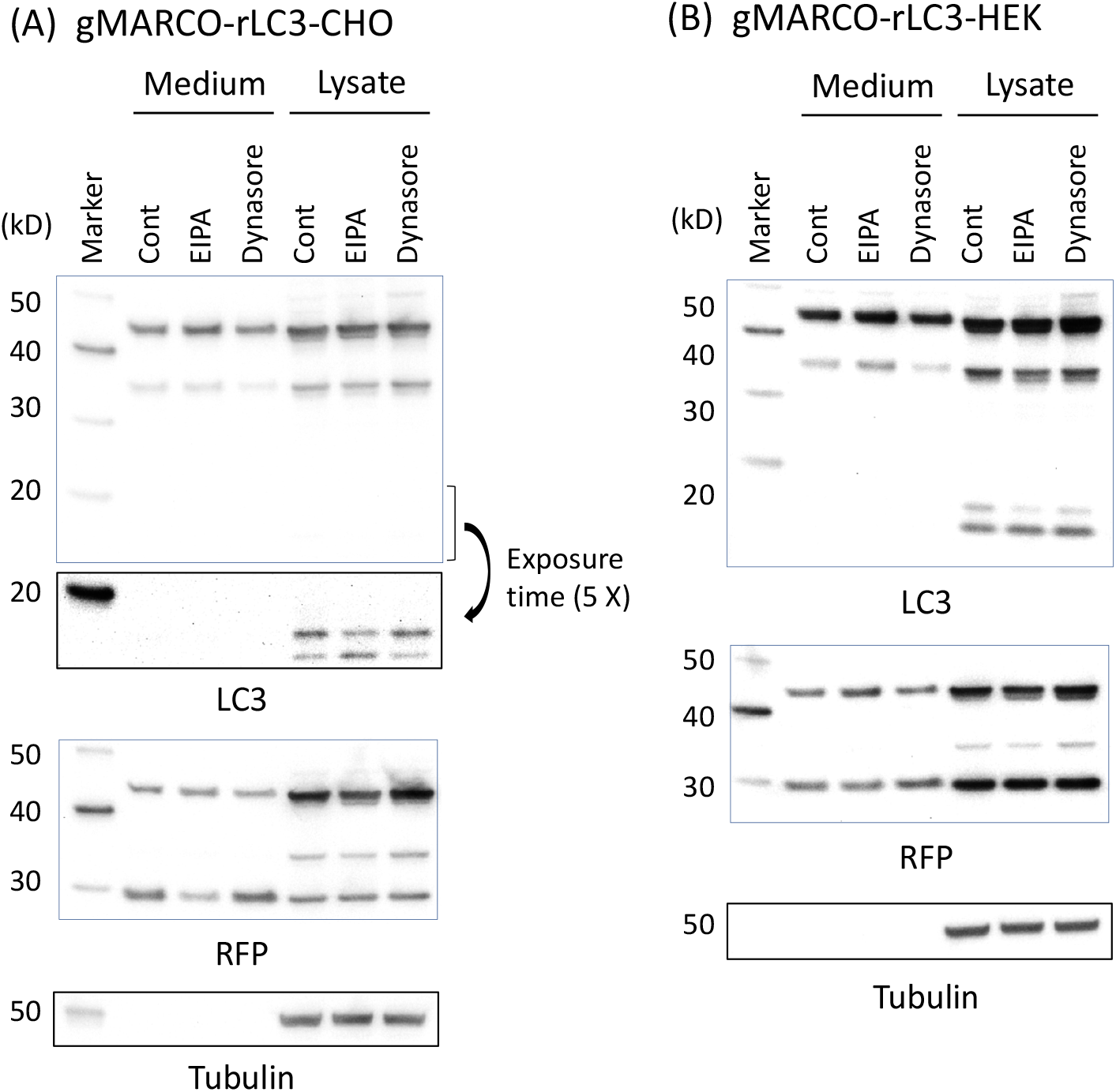
Western blot analyses of RFP-LC3 in the culture supernatant and cell lysate in gMARCO-rLC3-CHO (A) and gMARCO-rLC3-HEK cells (B). The early confluent cells were cultured in FBS-free Opti-MEM I medium in the presence of 0.2% DMSO (Cont), 50 μM EIPA (an inhibitor of macropinocytosis), and 50 μM dynasore (an inhibitor of endocytosis) for 16 h in 60 mm culture dishes. The cell-free conditioned medium was concentrated by 50 folds by ultrafiltration. The cells were lyzed with 120 μL RIPA buffer on ice for 10 min and the lysate was centrifuged at 9000 × g for 5 min. LC3, RFP, and tubulin in both the concentrated conditioned medium and the supernatant of cell lysates were analyzed by western blotting.

## Discussion

### Adsorption of proteins on the particle surface

The protein adsorption on the silica and polystyrene nanoparticles occurs in 0.5 min when the particle are exposed to human plasma, and fingerprints of particle-associated plasma proteins do not change qualitatively on prolonged exposure (Tenzer et al. 2013). Our present study indicated that cytosolic proteins were adsorbed on the surface of TiO_2_ particles even in the presence of 10% FBS (Fig.3 and Supplementary Fig. 2). In the conventional culture, the cell population changes according to the cell growth and cell death. In the normal turnover process, the cellular components are released in the culture medium. A recent secretosome study indicates that only 12.5% of the proteins detected in the culture medium of Huh7 cells contained signal peptides, suggesting that a large subset of proteins in the culture medium represents intracellular proteins released by injury of plasma membranes (Abbineni et al. 2022). However, the rapid release of the temperature-dependent release of cytosolic proteins in our current study indicated that cellular components were released into the culture medium from viable cells. The release of cytosolic proteins from gMARCO-rLC3-CHO cells was faster than that from gMARCO-rLC3-HEK cells (Figs. 4 and 6B). The cytoplasmic components including RFP-LC3 may have been released faster via the active movements of dendritic structures in gMARCO-rLC3-CHO cells (Supplementary Movie 1).

It has been reported that BSA is adsorbed on ZnO nanoparticles quickly just after preparation of the suspension, and the helix-dominant secondary structure of BSA is converted to more disordered structure (Ciobanu et al. 2022). Proteins adsorbed on the particle surface endow new biological identities, and change cellular internalization pathway and the following toxicological outcome (Ge et al. 2015). The hemolytic activity and cytotoxic effects of nano-silica were significantly reduced by blood plasma proteins formed on the silica particles (Tenzer et al. 2013). Bio-corona formed on nanoparticles plays a role in innate and adaptive immunity, and has been implicated in nanodrug delivery development and nanomedicine (Neagu et al. 2017). Our current results underscore the importance of cytosolic proteins released from the cultured cells in the recognition of particles by the cell membrane.

### Recognition of the particle surface by MARCO and the toxicological consequences

Phagocytosis is an important mechanism for clearance of environmental particles deposited in the respiratory region. The inefficient clearance of nanoparticles by alveolar macrophages leads to translocation of the particles into the lung tissue and potentially enhances adverse health effects (Geiser et al. 2008). MARCO, a plasma membrane receptor of phagocytes, plays a role in the clearance of environmental particles (Kobzik 1995). It has been reported that expression of MARCO in several types of cells results in morphological changes including formation of lamellipodia-like structures and of long dendritic processes (Kimura et al. 2007). The segment of the cysteine-rich domain V is important for the morphoregulatory activity of MARCO (Kimura et al. 2007) as well as TLR2-mediated inflammatory responses of macrophages (Novakowski et al. 2016). Our current study indicated that the dendritic structures of gMARCO-rLC3-CHO cells actively extended, retracted, and swelled along with the movement of cytosolic components (Supplementary Movie 1). It is plausible that MARCO on the dendritic structures recognized proteins adsorbed on the plastic dish bottom and was responsible for the movement of the dendritic structures.

*In vitro* and *in cellulo* toxicity assays have been widely used for safety screening of materials before they are handled in the market. There are several factors to determine the toxicity of particles such as solubility, size, shape, aspect ratio, chemical composition, and surface charge (Hirano 2009). Fibrous particles were more cytotoxic than spherical ones probably the integrity of plasma and lysosomal membrane is more easily damaged by fibrous bio-persistent particles (Hirano et al. 2000; Hirano et al. 2008). It has been shown that NH_2_-modified polystyrene 100 nm nanoparticles, but not carboxyl-or modified particles of the same particle size, were cytotoxic for human peripheral blood monocyte-derived macrophages. The NH_2_-modified polystyrene nanoparticles induced lysosomal damage late after 72 h via “proton sponge” effect, and triggered NLRP3 inflammasome activation resulting in the release of proinflammatory interleukin 1β (IL-1β) (Lunov et al. 2011b). Our present study suggests that the surface charge or zeta-potential of the particles can be modified by the protein corona formed on the particle surface in *in vitro/cellulo* toxicity assay. Information on the interaction of nanoparticles with biomolecules/cell membranes is prerequisite for designing engineered nanoparticles with enhanced or suppressed cellular uptake (Shang et al. 2014). Therefore, the adsorption of proteins on the particle surface should be carefully investigated for the safety evaluation of nanomaterials.

In summary, the surface of TiO_2_ particles adsorbs cellular proteins quickly irrespective of the presence of FBS in the cell culture system. Accordingly, TiO_2_ particles were coated with cytosolic proteins before association with MARCO on the plasma membrane of CHO cells. The foreign particulate substances of various components are recognized by MARCO. Thus, it is reasonable to conclude that MARCO, which functions as an adhesion receptor for environmental particles, recognizes intracellular proteins adsorbed on the particle surface rather than the “bare” surface of particles.

## Supporting information

This article contains supporting information.

## Author contributions

SH designed the study, performed the experiments, and drafted the paper. SK performed the experiments partially and revised the paper.

