## Supplementary figures and legends for "Macrophage Receptor with collagenous structure (MARCO) recognizes the cellular protein corona formed on environmental particles"

Supplementary figure legends

**Supplementary Fig. 1.** GFP-MARCO-CHO cells were loaded with 50 μg/mL pHrode red dextran for 15 hr. The cells were washed 3 times with HBSS. The cells were further cultured in the presence (ChQ) or absence (Cont) of 50 μM chloroquine for 8 hr. Note that red fluorescence of pHrode red dextran disappeared and green fluorescence of GFP-MARCO became brighter in lysosomes after alkalinization with chloroquine.

**Supplementary Fig. 2.** The conditioned medium was obtained after sub-culture of gMARCO-rLC3-CHO cells for three days in complete F12 culture medium containing 10% FBS. S-TiO2 or f-TiO2 particles were incubated at room temperature for 1 h in 4 mL of fresh culture medium or the supernatant of conditioned medium at a concentration of 10 μg/mL. The particles were collected by centrifugation at 9000 × g for 5 min and washed 2 times with PBS. Proteins adsorbed on the surface of the particles were extracted by heating (98 ℃) the particles with 40 μL of 1× SDS sample buffer for 3 min. A 10 μL aliquot was loaded to the gel.

**Supplementary Movie 1. The Live cell image of gMARCO-rLC3-CHO cells.** The cells were cultured in complete culture medium in a glass-bottom culture dish and exposed to 10 μg/mL f-$TiO_2$ for 1 h. Confocal microscopic frames for green and red fluorescence and bright field (*x-y-t* mode) were recorded every 5 min. The movie was built up at a rate of 3 frames per second for the overlaid frames. An arrow indicates the site where the cytoplasm was squeezed through the dendritic structures like "pressurized cytoplasmic streaming."

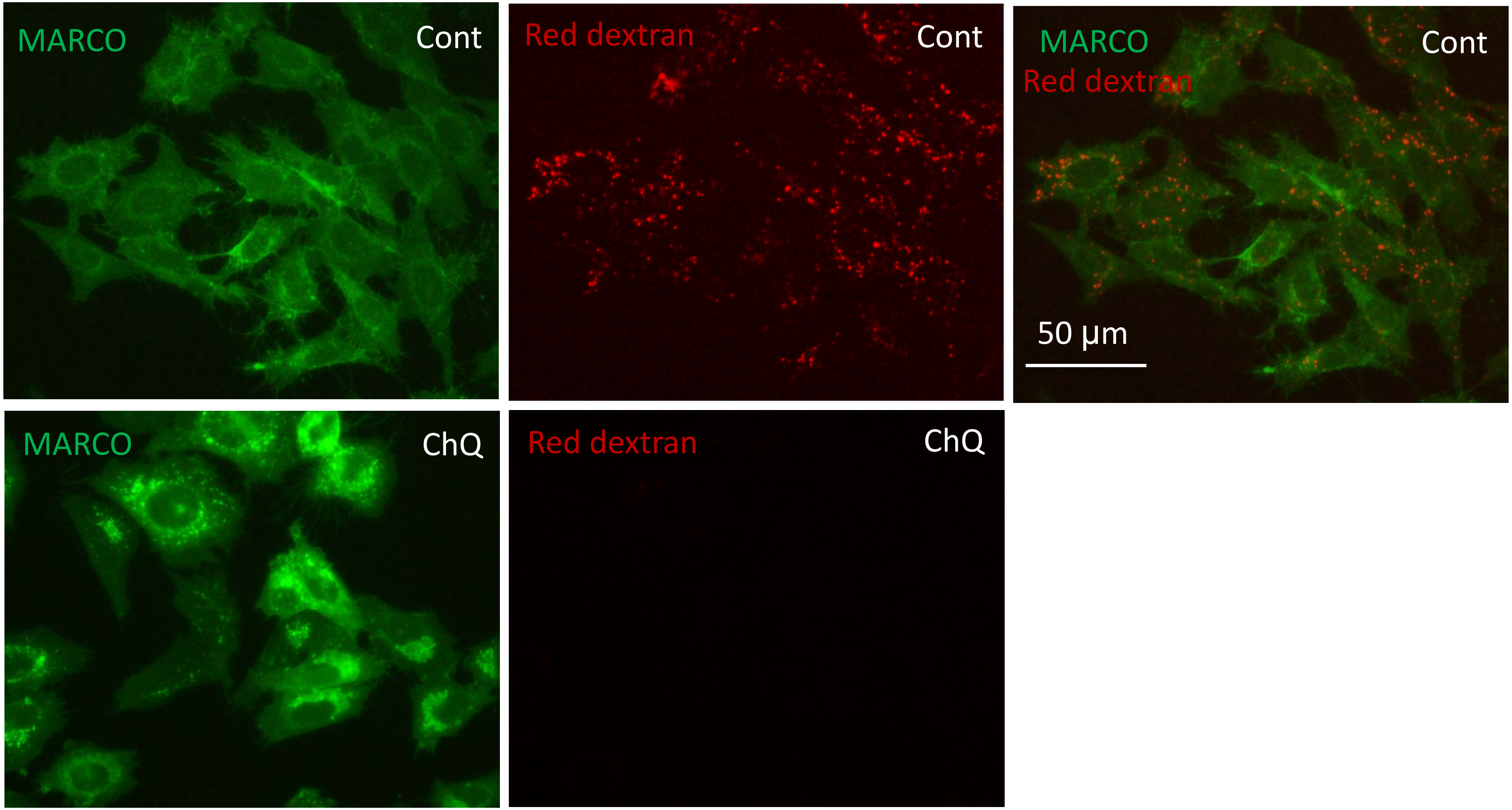


Supplementary Fig. 1

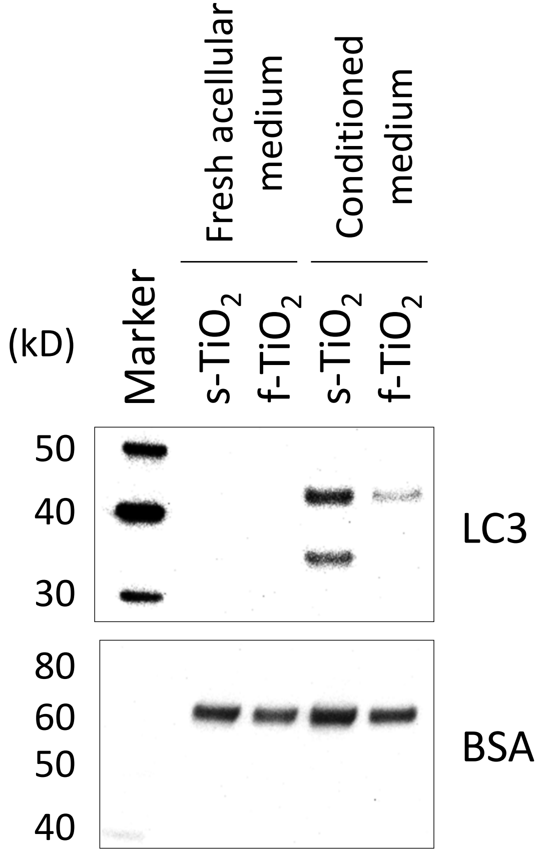


Supplementary Fig. 2
